# Impact of Reduced Spectral Resolution on Temporal-Coherence-Based Source Segregation

**DOI:** 10.1101/2024.03.11.584489

**Authors:** Vibha Viswanathan, Michael G. Heinz, Barbara G. Shinn-Cunningham

**Affiliations:** Neuroscience Institute, Carnegie Mellon University, Pitttsburgh, PA 15213; Department of Speech, Language, and Hearing Sciences, Purdue University, West Lafayette, IN 47907

## Abstract

Hearing-impaired listeners struggle to understand speech in noise, even when using cochlear implants (CIs) or hearing aids. Successful listening in noisy environments depends on the brain’s ability to organize a mixture of sound sources into distinct perceptual streams (i.e., source segregation). In normal-hearing listeners, temporal coherence of sound fluctuations across frequency channels supports this process by promoting grouping of elements belonging to a single acoustic source. We hypothesized that reduced spectral resolution—a hallmark of both electric/CI (from current spread) and acoustic (from broadened tuning) hearing with sensorineural hearing loss—degrades segregation based on temporal coherence. This is because reduced frequency resolution decreases the likelihood that a single sound source dominates the activity driving any specific channel; concomitantly, it increases the correlation in activity across channels. Consistent with our hypothesis, predictions from a physiologically plausible model of temporal-coherence-based segregation suggest that CI current spread reduces comodulation masking release (CMR; a correlate of temporal-coherence processing) and speech intelligibility in noise. These predictions are consistent with our behavioral data with simulated CI listening. Our model also predicts smaller CMR with increasing levels of outer-hair-cell damage. These results suggest that reduced spectral resolution relative to normal hearing impairs temporal-coherence-based segregation and speech-in-noise outcomes.

## 2. Introduction

Even when using state-of-the-art hearing aids and cochlear implants (CIs), people with sensorineural hearing loss (SNHL) find it significantly harder to understand speech in background noise than do listeners with clinically normal hearing (Hochberg et al., 1992; Dorman et al., 1998; Zeng, 2004; Chung, 2004; McCormack and Fortnum, 2013; Lesica, 2018). Prior studies suggest that reduced spectral resolution in electric/CI hearing (e.g., from current spread; Liang et al., 1999; Stickney et al., 2006) and in acoustic hearing with SNHL [e.g., from broadened tuning due to outer-hair-cell (OHC) damage; Sellick et al., 1982; Festen & Plomp, 1983] may contribute to the speech-in-noise deficits observed in hearing-impaired populations (Hall et al., 1988; Ter Keurs et al., 1992, 1993; Baer and Moore, 1993, 1994; Fu et al., 1998; Nelson et al., 2003; Stickney et al., 2004; Fu and Nogaki, 2005; Oxenham and Kreft, 2014). Decreased spectral resolution increases energetic masking within each frequency channel. Moreover, it may also impair the brain’s ability to perceptually separate different sound sources in an acoustic mixture (source segregation), like speech from noise. Although intact peripheral hearing and frequency resolution are posited as important for segregating speech from background noise, the neurophysiological mechanisms by which source segregation may fail in hearing-impaired populations is still poorly understood.

Manipulating a masker’s modulation spectrum to be less similar to that of the target sound reduces modulation masking (masking of target modulations or envelopes in a modulation-frequency-specific manner) in listeners with normal hearing (Bacon and Grantham, 1989; Stone and Moore, 2014; Viswanathan et al., 2021a). However, both CI users (Nelson et al., 2003; Stickney et al., 2004; Cullington and Zeng, 2008) and hearing-impaired listeners relying on acoustic hearing (Festen and Plomp, 1990; Bacon et al., 1998; Hall et al., 2012) show little to no release from modulation masking when the masker’s modulation spectrum is altered to be less like that of the target, an observation that is in line with the possibility that reduced spectral resolution interferes with source segregation. Prior studies, while describing the phenomenon, do not establish the mechanism explaining why reduced spectral resolution increases modulation masking.

According to the temporal-coherence theory of auditory scene analysis, temporally coherent sound modulations help group together sound elements from distinct frequency channels to form a perceptual object, thereby aiding segregation or unmasking of a target sound source from other competing sources (Elhilali et al., 2009; Teki et al., 2013; Viswanathan et al., 2021a, 2022). As a consequence, masker components that are temporally coherent with the target but in distinct frequency channels not driven by the target may interfere with target encoding and perception.

Compared to listeners with normal hearing, CI users (Ihlefeld et al., 2012; Zirn et al., 2013; Pierzycki and Seeber, 2014) and listeners with SNHL who rely on acoustic hearing (Hall et al., 1988; Moore et al., 1993; Ernst et al., 2010) show reduced comodulation masking release (CMR), a correlate of across-channel temporal-coherence-based segregation. Decreased frequency selectivity has been suggested as an explanation for this reduction in CMR in hearing-impaired individuals (Hall et al., 1988; Grose and Hall, 1996). Based on these prior results, we hypothesized that reduced spectral resolution, which occurs mainly due to current spread in electric hearing with CIs and due to OHC damage in acoustic hearing with SNHL, would adversely impact across-channel temporal-coherence-based source segregation and in turn speech understanding in noise. Specifically, decreased spectral resolution should increase the correlation between activity in distinct frequency channels by increasing target-masker overlap within each channel (i.e., reducing the sparsity of target and masker representations; Swaminathan and Heinz, 2011). We posited that these representational changes, jumbling together the neural responses to distinct, uncorrelated physical sources and increasing the temporal correlation of different channels, would disrupt source segregation and decrease CMR.

To test our hypothesis, we used a combination of physiologically plausible computational modeling and behavioral experiments. We based our approach on a wideband-inhibition-based model of across-channel temporal-coherence processing [developed to explain cochlear nucleus (CN) CMR data; Pressnitzer et al., 2001; expanded to predict speech confusions in different listening conditions; Viswanathan et al., 2022]. We compared model predictions of CMR and speech intelligibility in noise as a function of CI vocoding and current spread to behavioral measurements with simulated CI listening. We also obtained predictions for CMR as a function of degree of simulated OHC damage.

## 3. Materials and Methods

### 3.1. Stimulus generation

#### 3.1.1. CMR stimuli to evaluate temporal-coherence processing

Figure 1 illustrates the stimuli used to model and behaviorally measure CMR. The stimuli consisted of a 3022-Hz tone (the target signal) in a sinusoidally amplitude-modulated (SAM) tonal complex masker. The masker was composed of three SAM tones, at carrier frequencies of 3022 Hz (on-frequency component; OFC), 2142 Hz (first flanking component), and 4264 Hz (second flanking component). Note that these target and masker frequencies were chosen so as to align with the filters used during vocoding (see section 3.1.4.). Each of the flankers was separated from the OFC (and the target signal) by three times the equivalent rectangular bandwidth (ERB) of the psychophysical tuning curve at the target-signal frequency for normal-hearing listeners (Glasberg and Moore, 1990). A 10 Hz modulation rate and 100% modulation depth were used for all of the SAM tones. The two flankers were each presented at the same sound level as the OFC.

**Figure 1.**
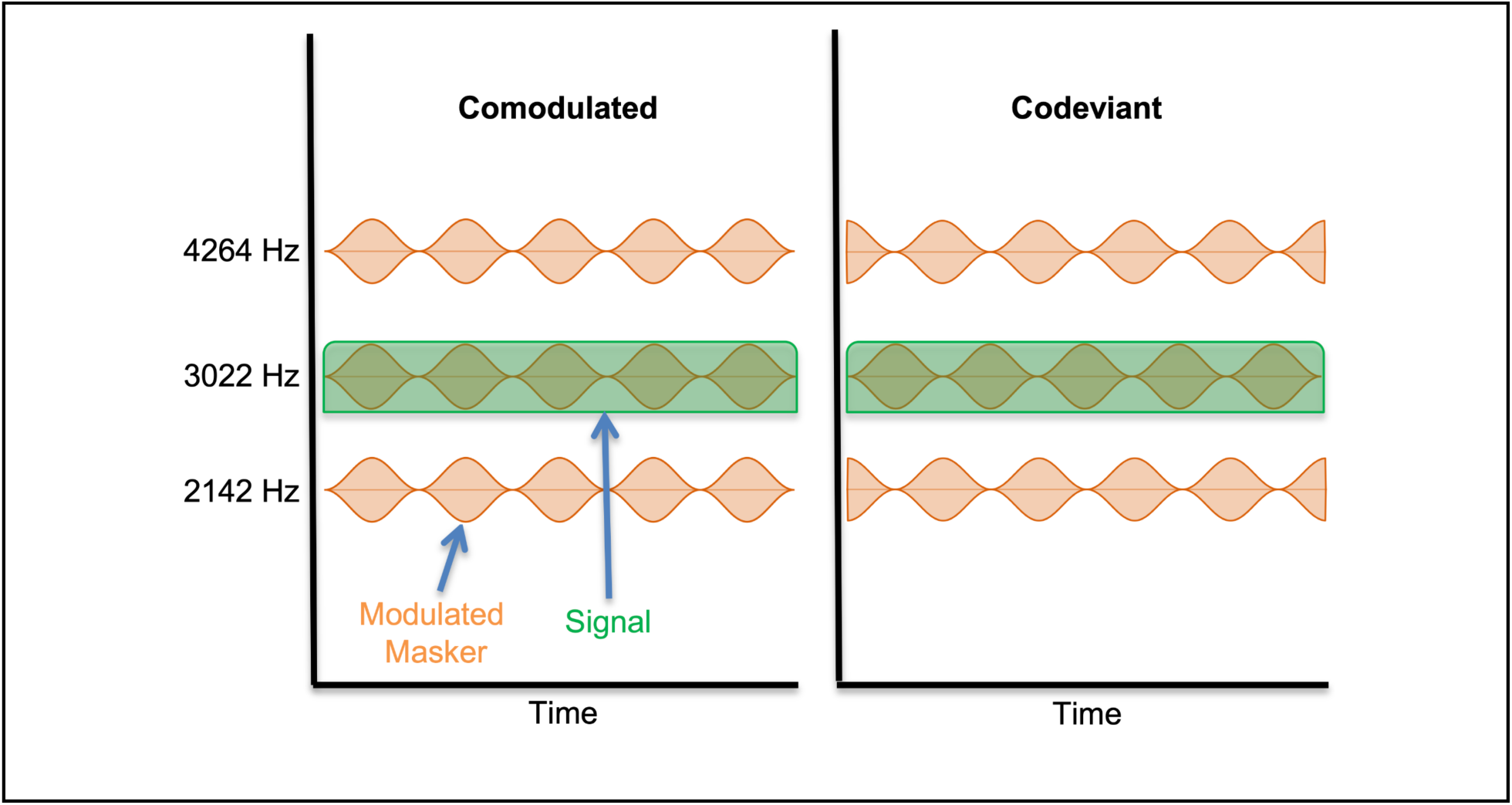
Comodulated making release (CMR) stimuli used for computational modeling and behavioral measurements. The stimuli consisted of a target signal (shown in green) in a 100% sinusoidally amplitude-modulated (SAM) tonal complex masker (shown in orange). The masker was composed of three 10-Hz SAM tones at carrier frequencies of 3022 Hz (on-frequency component; OFC), 2142 Hz (first flanking component), and 4264 Hz (second flanking component). All three SAM tones were presented at the same sound level. In the Comodulated condition, the flanking components were modulated in phase with the OFC, while in the Codeviant condition, they were modulated 180° out of phase with the OFC. The target signal was a 3022 Hz pure tone presented at different signal-to-noise ratios (SNRs). The level of the OFC was fixed while that of the target signal was varied according to the SNR. The total duration of each stimulus was 0.5 seconds.

Stimuli were created for two CMR conditions: (i) In the Comodulated (temporally coherent) condition, the flanking components were modulated in phase with the OFC, and (ii) In the Codeviant condition, the flankers were modulated 180° out of phase with the OFC. In each condition, the target signal was presented at different signal-to-noise ratios (SNRs; defined as the ratio of target-signal power to OFC power). For computational modeling, we used SNRs of 12, 6, 0, -6, -12, -18, and -inf (corresponding to no signal being presented) dB for both CMR conditions. For the behavioral experiment, we used SNRs of 6, 0, -6, -12, -18, and -24 dB for the Comodulated condition, and SNRs of 12, 6, 0, -6, -12, and -18 dB for the Codeviant condition. For both computational modeling and the behavioral experiment, the root mean square value (RMS) of the OFC was fixed while that of the target signal was varied according to the SNR. The total duration of each stimulus was 0.5 seconds.

#### 3.1.2. Consonant identification stimuli

The stimuli used to model and behaviorally measure consonant identification in noise consisted of twenty consonants from the Speech Test Video (STeVi) corpus (Sensimetrics), namely /b/, /tʃ/, /d/, /ð/, /f/, /g/, /dʒ/, /k/, /l/, /m/, /n/, /p/, /r/, /s/, /ʃ/, /t/, /θ/, /v/, /z/, and /ʒ/. The consonants were presented in consonant-vowel (CV) context, where the vowel was /a/. Two tokens of each CV were included, one spoken by a female and one by a male talker, to reflect real-life talker variability. The CV utterances were embedded in the carrier phrase, “You will mark /CV/ please”, to create natural running speech. Stimuli were created for (i) speech in quiet (SiQuiet), and (ii) speech in speech-shaped stationary noise (SiSSN) masking conditions. To create SiSSN, speech was added to stationary Gaussian noise at -2 dB SNR (SNR chosen by piloting to yield relatively high speech intelligibility for intact stimuli, which minimized the likelihood of behavioral floor effects for cochlear-implant-processed stimuli) such that the masking noise started 1 second before the target speech and continued for the entire duration of the trial; this was done to cue subjects’ attention to the stimulus before the target sentence was played. The long-term spectra of the target speech (including the carrier phrase) and that of the stationary noise were matched. The RMS of the target speech was set to a fixed value across all of the consonant identification stimuli.

#### 3.1.3. Stimuli used for online volume setting

In the online CMR and speech identification experiments, listeners were asked to set the stimulus levels to be comfortably loud. The stimuli used for this volume setting were specific to the experiment. In both cases, the stimulus presented during volume-setting was relatively long to ensure listeners had sufficient time to settle on an appropriate level (see Section 3.3.2 for the specific instructions given to the listeners).

The volume-set stimulus used in the CMR experiment consisted of six Comodulated stimuli at the different SNRs used in the actual experiment stitched together to obtain a stimulus with a total duration of ∼30 seconds. By including all of the SNR conditions in the volume-set stimulus (even though the overall sound levels differ across SNRs), this approach ensured that listeners are comfortable with the volume for all SNR conditions used in the experiment.

The volume-set stimulus used in the consonant identification experiment was created by stitching together 15 speech sentences [from the Harvard/Institute of Electrical and Electronics Engineers lists (Rothauser, 1969), spoken in a female voice and recorded as part of the PN/NC corpus (McCloy et al., 2013)] mixed with speech-shaped noise at -2 dB SNR. The total duration of this stimulus was ∼one minute. The RMS of the target speech was set to be equal to that of the target speech in the main consonant identification experiment.

#### 3.1.4. Cochlear-implant processing

To explore the role of spectral smearing in CI listening, we processed CMR and consonant identification stimuli with three different levels of CI simulation (hereafter referred to as the three vocoding conditions): (i) Intact, (ii) Vocoded, and (iii) Current Spread. The Intact stimuli are described in Sections 3.1.1. and 3.1.2. To create stimuli for the Vocoded condition, Intact stimuli were subjected to cochlear-implant processing. Specifically, subband signals were extracted by band-pass filtering (using a sixth-order Butterworth filter) Intact stimuli at vocoder center frequencies and channel cutoffs matching those used in Advanced Bionics CIs (Table 1; 16 vocoder channels in total, spanning frequencies between 250 and 8700 Hz). The envelope in each subband was extracted by half-wave rectifying and low-pass filtering (with a sixth order Butterworth filter) the subband signal up to a maximum of 5% of the center frequency (to avoid artifacts that may be produced if the envelope frequency gets resolved at an individual listener’s cochlea). Then, the extracted envelopes were used to modulate pure-tone carriers at the corresponding center frequencies (Table 1). Results were summed across carrier bands to generate the final stimuli. To create stimuli for the Current Spread condition, the same procedure as above was used but with the extra step of smearing the extracted envelopes across frequency channels before modulating the pure-tone carriers. Specifically, a current spread of 8 dB per octave was simulated via the spectral smearing operation described in Equation 1 (following Nelson et al., 2011, and Oxenham and Kreft, 2014).

**Table 1.** Center frequencies and cutoffs (high, low; in Hz) for the vocoder channels in Advanced Bionics’ cochlear implants (CIs).

| High | Center | Low |
| --- | --- | --- |
| 416 | 333 | 250 |
| 494 | 455 | 416 |
| 587 | 540 | 494 |
| 697 | 642 | 587 |
| 828 | 762 | 697 |
| 983 | 906 | 828 |
| 1168 | 1076 | 983 |
| 1387 | 1278 | 1168 |
| 1648 | 1518 | 1387 |
| 1958 | 1803 | 1648 |
| 2326 | 2142 | 1958 |
| 2762 | 2544 | 2326 |
| 3281 | 3022 | 2762 |
| 3898 | 3590 | 3281 |
| 4630 | 4264 | 3898 |
| 8700 | 6665 | 4630 |

Let *e_i_* be the original temporal envelope extracted in subband *i*. Then *E_i_*, the envelope after spectral smearing, is calculated as a function of time *t* as

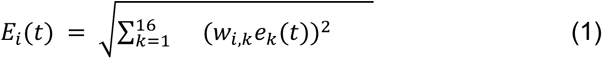

where *w_i,k_* is the weight applied to *e_k_*(*t*) to derive the smeared envelope *E_i_*(*t*); this weight corresponds to an attenuation of 8 dB/octave on either side of subband *i*. Note that the RMS values of all stimuli were matched across the three vocoding conditions at each SNR.

### 3.2. Physiologically based computational modeling

We used an across-channel temporal-coherence-based source-segregation model (Figure 2) developed and validated in our prior work (Viswanathan et al., 2022) to predict CMR and speech intelligibility in noise as a function of simulated CI listening, and to predict CMR for Intact stimuli as a function of degree of OHC damage. The source-segregation model is described in detail in our prior work (Viswanathan et al., 2022) and hence only briefly reviewed below. The first stage of the source-segregation model simulates the auditory periphery using the Bruce et al. (2018) auditory-nerve (AN) model with the parameters described in Table 2. Since all of the stimuli used in this study contain the same audio signal across the left and right channels, the AN model was provided with only one (versus two) audio channel input. One hundred and fifty stimulus repetitions were used to derive peristimulus time histograms (PSTHs) from model auditory-nerve outputs with a PSTH bin width of 1 ms (i.e., 1 kHz sampling rate). Outputs from the AN model were input into a CMR circuit model (Figure 2C), which simulates across-channel temporal-coherence processing that mirrors computations in the ventral CN (Pressnitzer et al., 2001).

**Figure 2.**
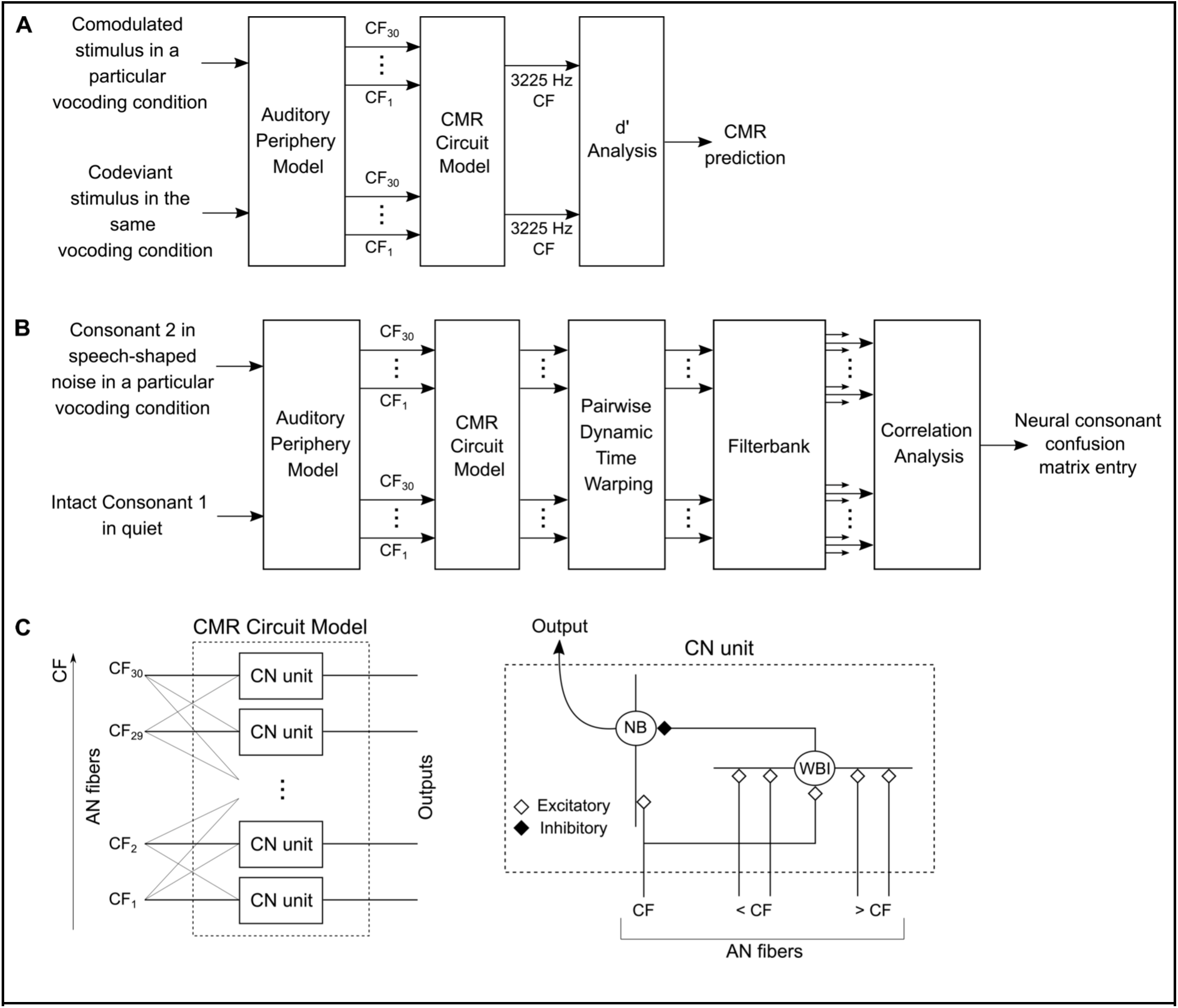
Across-channel temporal-coherence-based source-segregation model (Viswanathan et al., 2022) to predict CMR (Panel A) and consonant confusions in noise (Panel B) in different vocoding conditions (Intact, Vocoded, Current Spread) and as a function of outer-hair-cell (OHC) damage. Panel C illustrates the CMR circuit model shown in Panels A and B. Detailed descriptions and model parameters are provided in the main text.

**Table 2.** Parameters of the auditory-nerve (AN) model (Bruce et al., 2018) used in the current study.

| Name | Value |
| --- | --- |
| Number of cochlear filters | 30 |
| Characteristic frequencies (CFs) | Equally spaced on an equivalent rectangular bandwidth (ERB)-number scale (Glasberg and Moore, 1990) between 125 and 8000 Hz |
| Outer hair cell function ($C_{OHC}$) | To derive predictions for different vocoding conditions (Intact, Vocoded, Current Spread), this parameter was set to Normal; to derive predictions for different levels of OHC damage, this parameter was varied from 1 (Normal) to 0 (complete dysfunction) in logarithmic steps |
| Inner hair cell function ($C_{IHC}$) | Normal |
| Species and frequency tuning | Human with the Shera et al. (2002) cochlear tuning at low sound levels; with suppression, the Glasberg and Moore (1990) tuning is effectively obtained for our broadband, moderate-level stimuli (Heinz et al., 2002; Oxenham and Shera, 2003) |
| Noise type for inner-hair-cell synapse model | Fixed fractional Gaussian noise |
| Spontaneous firing rate | Medium (10 spikes/second) |
| Power-law adaptation dynamics in the synapse | Approximate implementation |
| Absolute refractory period | 0.6 ms |
| Relative refractory period | 0.6 ms |

CN units at different characteristic frequencies (CFs) form the building blocks of the CMR circuit model (Figure 2C). Each CN unit consists of a narrowband cell (NB) that receives narrow on-CF excitatory input from the AN and inhibitory input from a wideband inhibitor (WBI). The WBI receives excitatory inputs from AN fibers tuned to CFs spanning 2 octaves below to 1 octave above the CF of the NB that it inhibits. The time constants for the excitatory and inhibitory synapses are 5 ms and 1 ms, respectively. The WBI input to the NB is delayed with respect to the AN input by 2 ms. The excitation-to-inhibition ratio was set to 1.75:1. Note that all of the parameters of the CMR circuit model used in this study are exactly the same as in our prior work that validated the overall source-segregation model with these parameters (Viswanathan et al., 2022).

To determine stimulus levels for computational modeling, we generated model AN threshold tuning curves and rate-level curves for different degrees of OHC damage. All tuning and rate-level data were obtained for a 3225 Hz CF (i.e., the CF at which we derived CMR predictions; see Figure 2A).

To generate threshold tuning curves, we presented a 0.5-second-long tonal signal varying in frequency and level to the AN model. For each tone frequency, tone level, and degree of OHC damage, we computed the difference between the time-averaged firing rate in the steady-state portion of the model AN response (25 ms onwards) and the corresponding firing rate in the absence of the tonal signal. The tone detection level threshold corresponding to a firing-rate difference of 10 spikes/s (Liberman, 1978) was computed for the different tone frequencies and degrees of OHC damage. The resulting threshold tuning curves are shown in Figure 3A.

**Figure 3.**
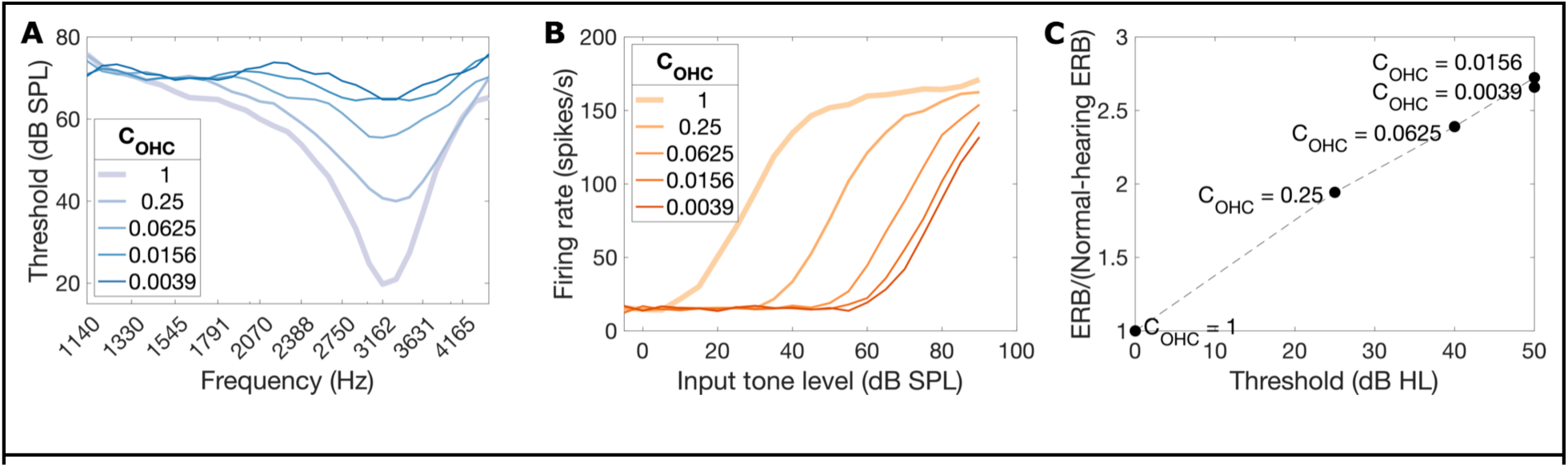
Model AN-fiber threshold tuning curves (Panel A), rate-level curves (Panel B), and thresholds (in dB HL, i.e., relative to C_OHC_ = 1) plotted against ERB [Panel C; ERB was derived from Panel A tuning data, and expressed as a ratio relative to the ERB of a normal-hearing (C_OHC_ = 1) ear]. Data are shown for varying degrees of OHC damage [simulated by varying the model parameter C_OHC_ from 1 (normal) to 0 (complete OHC dysfunction)]. All data were obtained for a 3225 Hz CF (i.e., the CF at which we derived CMR predictions).

To generate rate-level curves, we presented a 0.5-second-long 3225 Hz tone at various levels to the AN model. The time-averaged firing rate during the steady-state portion (25 ms onwards) of the model AN response was used to derive model AN rate-level curves for different degrees of OHC damage (Figure 3B).

Figure 3C shows the relationship between the simulated degree of OHC damage and ERBs derived from Figure 3A tuning data. The model AN-fiber data in Figure 3C are comparable to psychophysical frequency selectivity data obtained from individuals with cochlear hearing loss (Moore, 1996).

Figure 2A illustrates the steps used to predict CMR. The CMR circuit model was simulated at a 3225 Hz CF, which is the particular CF from Table 2 that is closest to the carrier frequency of the OFC (3022 Hz) in the CMR stimuli. To predict CMR in the different vocoding conditions (Intact, Vocoded, Current Spread), we used a fixed OFC level of 48 dB sound pressure level (SPL) for two reasons: (1) the 48 dB SPL OFC has the same energy within an ⅓-octave band as a 60 dB SPL conversational-level broadband sound (assuming pink spectrum spanning 250–8000 Hz), and (2) the normal-hearing model-AN pure-tone threshold at CF is ∼20 dB SPL (Figure 3A) and so at the worst stimulus SNR (-18 dB), the target would be at least 10 dB sensation level (SL) or 30 dB SPL. This choice of level yielded a firing rate at the output of the model AN that was greater than the spontaneous rate but that did not saturate in response to the loudest stimulus.

To predict CMR for Intact stimuli as a function of degree of OHC damage, we performed two different simulations: (1) The first used a fixed OFC level of 83 dB SPL to ensure that the target signal would be “audible”, i.e., generate model AN responses greater than spontaneous rate, for even the greatest degree of OHC damage (for which the model-AN pure-tone threshold is ∼65 dB SPL; Figure 3A) and worst SNR condition (-18 dB), and (2) The second used a fixed loudness (versus fixed SPL) for the OFC. For this simulation, the OFC level for normal hearing was fixed at 48 dB SPL, as in the CI-listening simulation. To determine the OFC levels needed to achieve equal OFC loudness as a function of OHC damage, we used the predictions from the Moore and Glasberg (1998, 2004) loudness model.

For all of the above CMR predictions, we used the time-averaged statistics of the CMR circuit model output firing rate in the absence of the target signal to compute null distributions. For each vocoding/OHC-damage condition, stimulus repetition, CMR condition (Comodulated, Codeviant), and SNR, the time-averaged firing rate at the output of the CMR circuit model was compared with the corresponding null distribution to estimate the neurometric sensitivity, d’. CMR was calculated as the average SNR threshold difference between Codeviant and Comodulated conditions across the d’ values predicted by the model.

Figure 2B illustrates the steps used to predict speech intelligibility in the different vocoding conditions (approach established in our prior work; Viswanathan et al., 2022). The level for the target speech in the consonant identification stimuli was set to 60 dB SPL across all stimuli, i.e., a conversational level; this level produced sufficient (i.e., firing rate greater than spontaneous rate) model AN responses for consonants in quiet and also did not produce saturated responses to the loudest stimulus. AN model output PSTHs for the consonant identification stimuli were processed to retain only those time segments when the target consonants were presented. These segments were then input into the CMR circuit model. The CMR circuit model was simulated at the same set of CFs as the AN model (Table 2). Dynamic time warping was performed to align circuit model outputs across time for each pair of consonants. A filterbank comprising a low-pass filter with a 1 Hz cutoff in parallel with eight bandpass filters (octave spacing, quality factor of 1, and center frequencies between 2 and 256 Hz; Jørgensen et al., 2013) was used to decompose the warped outputs at each CF into different frequency bands. For each vocoding condition, consonant, talker, CF, and band, Pearson correlation coefficients were computed between the filterbank output for that consonant in speech-shaped noise and the output for each of all 20 consonants in quiet in the Intact condition (i.e., the output expected for a normal-hearing ear hearing the sounds in isolation). These correlations were squared, then averaged across talkers, CFs, and bands. Finally, for each modeled consonant (consonant 2), these average squared correlations were normalized such that their sum across all 20 consonants that could be reported (consonant 1) equaled one; this procedure yielded a neural consonant confusion matrix for each vocoding condition. The overall model (Figure 2B) was calibrated by fitting a logistic/sigmoid function mapping the model-derived neural consonant confusion matrix entries for the Intact SiSSN condition to corresponding perceptual measurements. The mapping derived from this calibration was used to predict perceptual speech intelligibility for SiSSN in the different vocoding conditions from the corresponding neural confusion matrices.

### 3.3. Behavioral experiments

#### 3.3.1. Participants

Data were collected on a web-based psychoacoustics platform (Mok et al., 2023) from anonymous subjects recruited using Prolific.co. The subject pool was restricted using a screening method developed by Mok et al. (2023), which contained three parts: (i) a survey that was used to restrict subjects based on age to 18–55 years (to exclude significant age-related hearing loss), whether or not they were US/Canada residents, US/Canada born, and native speakers of North American English (because North American speech stimuli were used), history of hearing and neurological diagnoses if any, and whether or not they had persistent tinnitus; (ii) headphone/earphone checks (hereafter referred to as headphone checks); and (iii) a speech-in-babble-based hearing screening. Subjects who passed the three-part screening were invited to participate in the CMR and consonant identification experiments, and when they returned, headphone checks were performed again. All subjects had completed at least 40 previous studies on Prolific and had >90% of these studies approved. These procedures were shown to successfully select participants with near-normal hearing status, attentive engagement, and stereo headphone use (Mok et al., 2023). Subjects provided informed consent in accordance with remote testing protocols approved by the Purdue University Institutional Review Board (IRB).

#### 3.3.2. Experimental design

We conducted two psychophysical experiments to predict the impact of CI vocoding and current spread on across-channel temporal-coherence-based source segregation; the first measured CMR and the second consonant identification in noise. Subjects performed the experiments using their personal computers and headphones (our online infrastructure included checks to prevent the use of mobile devices). All stimuli were presented diotically.

Headphone checks were performed at the beginning of each experiment using a paradigm validated by Mok et al. (2023). In this paradigm, subjects first performed a task that distinguishes between listening with a pair of free-field speakers versus using headphones (Woods et al., 2017). Subjects then performed a second task where the target cues were purely binaural, allowing us to test if headphones/earphones were used in both ears. This task was a three-interval three-alternatives-forced-choice task where the target interval contained white noise with interaural correlation fluctuating at 20 Hz, while the dummy intervals contained white noise with a constant interaural correlation. Subjects were asked to detect the interval with the most flutter or fluctuation. Only those subjects who scored greater than 65% in each of the two headphone-check tasks were allowed to proceed to the rest of the experiment.

Subjects performed a volume-adjustment task before each headphone check and also before the main task in each experiment. In the volume-adjustment task, subjects were asked to make sure that they were in a quiet room and wearing wired (not wireless) headphones or earphones, and not to use computer speakers. They were then asked to set their computer volume to 10%–20% of the full volume, after which they were played either a speech-in-babble stimulus (if the volume calibration was performed prior to headphone checks) or a volume-set stimulus more closely matched to the stimuli used in the actual experiment (see Section 3.1.3.). During this, they were asked to adjust their volume up to a comfortable, but not too loud level. Once subjects had adjusted their computer volume, they were instructed not to change the volume setting during the experiment to avoid sounds becoming too loud or soft.

For the CMR experiment, separate studies (two total) were posted on Prolific.co for the following two different orders of the vocoding conditions (“condition-orders”): (i) Intact, Vocoded, Current Spread, and (ii) Current Spread, Vocoded, Intact. Note that to reduce task confusion, we did not interleave the different vocoding conditions. Each study presented eight repetitions of each vocoding condition, CMR condition (Comodulated, Codeviant), and SNR. Ten subjects were used per CMR study (subject overlap between studies was not controlled). Thus, there were 20 combinations of subject and condition-order (i.e., N=20 samples total) in the CMR experiment. Within each study (i.e., a particular condition-order), all subjects performed the task with the same stimuli. All condition effect contrasts were computed on a within-subject basis and averaged across subjects. A four-alternatives-forced-choice (4-AFC) design was used. Subjects were instructed that in each trial they would hear four sounds and were asked to choose which of the four contained a steady beep. To promote engagement with the task, subjects received feedback after every trial as to whether or not their response was correct. Subjects were not told what the correct answer was to avoid over-training to the acoustics of the stimuli across the different vocoding conditions.

For the consonant identification experiment, separate studies (four total) were posted on Prolific.co for the two different talkers and the following two different orders of the vocoding conditions (condition-orders): (i) Intact, Vocoded, Current Spread, and (ii) Current Spread, Vocoded, Intact. Each of the four studies presented, in random order, one stimulus repetition per consonant in each vocoding condition. Twelve subjects were used per consonant identification study and subject overlap between studies was not controlled. Thus, there were a total of 48 combinations of subject, talker, and condition-order (i.e., N=48 samples total) in the consonant identification experiment. Within each study (a particular talker and condition-order), all subjects performed the task with the same stimuli. Moreover, all condition-effect contrasts were computed on a within-subject basis and then averaged across subjects. We chose stationary noise (versus babble) as the masking noise to minimize any masker instance effects (Zaar and Dau, 2015; Viswanathan et al., 2021b). We used different masker instances (i.e., realizations of stationary noise) for the different consonants and talkers; however, we did not vary the masker instance across the different vocoding conditions.

In each consonant identification study, just prior to the main consonant identification task, subjects performed a short demonstration (“demo”) task, which familiarized them with the overall consonant identification paradigm and with how each consonant sounds for the particular talker used in the study. Subjects were instructed that in each trial they would hear a voice say “You will mark <SOMETHING> please.” They were told that at the end of the trial, they would be given a set of options for <SOMETHING> and that they would have to click on the corresponding option. Consonants were first presented in quiet (SiQuiet) in sequential order from /b/ to /ʒ/. This order was matched in the consonant options shown on the screen at the end of the trial. After the stimulus ended in each trial, subjects were asked to click on the consonant they heard. After subjects had heard all consonants sequentially in quiet, they were tasked with identifying consonants presented in random order and spanning the same set of listening conditions as the main task in the experiment. Subjects were instructed to ignore any background noise and only listen to the voice saying, “You will mark <SOMETHING> please.” Only subjects who scored at least 85% in the demo’s SiQuiet condition were selected for the next stage of the experiment, so as to ensure that all subjects were able to perform the task. In the main task of the consonant identification experiment, subjects were given similar instructions as in the demo but told to expect trials with background noise from the beginning. As in the CMR experiment, here too subjects received feedback after every trial as to whether or not their response was correct to promote engagement with the task. Subjects were not told what consonant was presented to avoid over-training to the acoustics of the stimuli across the different vocoding conditions, except for the first part of the demo where subjects heard all consonants in quiet in sequential order.

### 3.4. Statistical analysis

To test for significant differences between Vocoded and Current Spread conditions in the behavioral CMR measurements, we used a linear mixed-effects model. Measured CMR served as the response, and vocoding condition (factor variable with three levels; Intact, Vocoded, and Current Spread) and sample (factor variable with N=20 levels, corresponding to 20 combinations of subject and condition-order) served as predictors. Vocoding condition was treated as a fixed-effects predictor and sample as a random-effects predictor. Anova (Type II Wald F tests with Kenward-Roger degree of freedom; Kenward and Roger, 1997) was used for statistical testing.

To test whether there are significant differences between Vocoded and Current Spread conditions in the behavioral speech-intelligibility-in-noise measurements, we used a linear mixed-effects model. Percent consonants correct in noise served as the response, and vocoding condition (factor variable with three levels; Intact, Vocoded, and Current Spread) and sample (factor variable with N=48 levels, corresponding to 48 combinations of subject, talker, and condition-order) served as predictors. Vocoding condition was treated as a fixed-effects predictor and sample as a random-effects predictor. Anova (Type II Wald F tests with Kenward-Roger degree of freedom; Kenward and Roger, 1997) was used for statistical testing.

To test whether the across-channel temporal-coherence model better predicts behavioral scores for percent consonants correct compared to the within-channel model, we computed the mean squared error between model predictions and behavioral data across stimulus repetitions and vocoding conditions. We used a nonparametric permutation-based approach (Nichols and Holmes, 2002) to generate realizations of the distribution under the null hypothesis that the mean squared error is the same for the across- and within-channel models. Specifically, to generate each realization, we randomly flipped the sign of the difference between the two models in the squared error (computed between model predictions and behavioral data) for each stimulus repetition and vocoding condition; then we computed the mean of the result across stimulus repetitions and vocoding conditions. In this way, we generated 100,000 realizations of the null distribution. Finally, the difference in the mean squared error between the two models with the correctly labeled data was compared with the null distribution to generate a p-value.

### 3.5. Code accessibility

Subjects were directed from Prolific.co to the SNAPlabonline psychoacoustics infrastructure (Bharadwaj, 2021; Mok et al., 2023) to perform the study. Offline data analyses were performed using custom software in MATLAB (The MathWorks, Inc., Natick, MA) and PYTHON (Python Software Foundation, Wilmington, DE). Statistical analyses were performed using R (R Core Team; www.R-project.org). Visualizations used the colorblind-friendly Colorbrewer (Harrower and Brewer, 2003) colormap palettes. The code for our computational model was published on GitHub at https://github.com/vibhaviswana/ modeling-consonant-confusions as part of our prior work (Viswanathan et al., 2022).

## 4. Results

### 4.1. Simulated CI current spread degrades temporal-coherence processing

Figures 4A,B show d’ estimates and CMR predictions at the output of the across-channel temporal-coherence-based source-segregation model as a function of simulated CI vocoding and current spread. CMR was predicted as the mean SNR threshold difference between Codeviant and Comodulated conditions across the d’ values predicted by the model. The temporal-coherence-based segregation model predicts a smaller CMR in the Current Spread condition compared to the Intact and Vocoded conditions. Figures 4C,D show behavioral measurements for proportion trials correct and CMR (N=20) in the same vocoding conditions. Behavioral CMR was calculated as the SNR threshold difference between Codeviant and Comodulated conditions at a percent-correct score of 66%. Behavioral measurements are consistent with model predictions and show statistically significant differences in CMR between Vocoded and Current Spread conditions [F(2,38) = 12.479, p = 6.82e-05]. These data support our hypothesis that current spread (and the resulting reduction in spectral resolution) in CIs degrades across-channel temporal-coherence-based segregation of a target sound source from background noise (of which CMR is a correlate).

**Figure 4.**
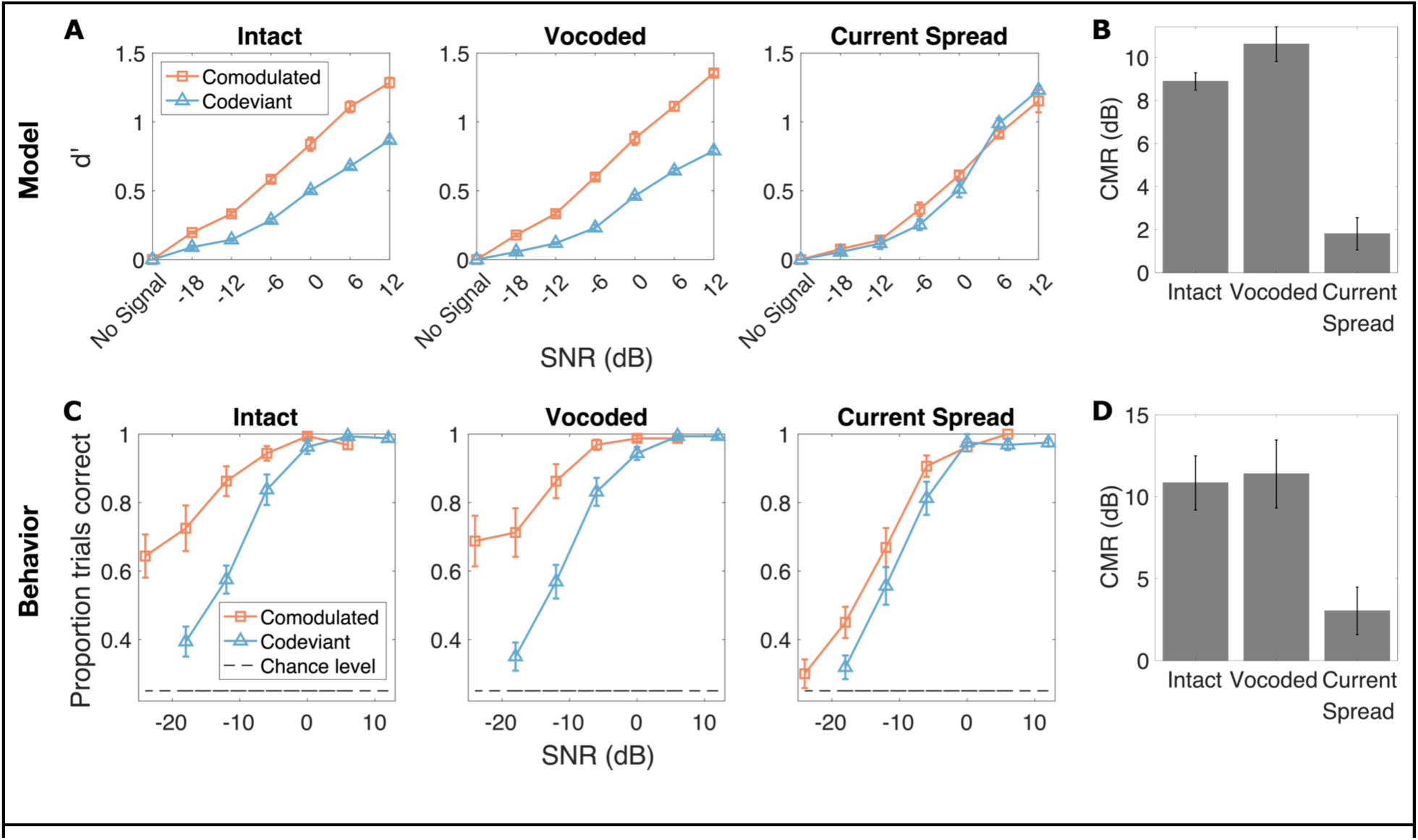
CMR as a function of simulated CI vocoding and current spread. Panel A shows estimated d’ values (mean and standard error across stimulus repetitions) from the across-channel temporal-coherence-based source-segregation model (Figure 2A) for different SNRs and CMR conditions (Comodulated, Codeviant). Panel B shows CMR predictions from the temporal-coherence model (mean and standard error across stimulus repetitions), calculated from Panel A as the mean difference in SNR threshold between Codeviant and Comodulated conditions across the d’ values predicted by the model. Panel C shows behaviorally measured proportion trials correct (mean and standard error across N=20 samples) for different SNRs and CMR conditions. Panel D shows behaviorally measured CMR (mean and standard error across N=20 samples), which was calculated for each sample as the SNR threshold difference between Codeviant and Comodulated conditions at a percent-correct score of 66%.

### 4.2. Simulated CI listening degrades speech-in-noise outcomes

Figure 5 shows model predictions (leftmost and middle plots) and behavioral measurements (rightmost plot; N=48) for percent consonants correct in speech-shaped noise as a function of simulated CI vocoding and current spread. Behavioral data show significant differences in percent consonants correct between Intact, Vocoded, and Current Spread conditions [F(2,94) = 318.87, p < 2.2e-16], suggesting that CI processing impacts speech-in-noise outcomes. The across-channel temporal-coherence-based model better predicts these behavioral outcomes across conditions than a purely within-channel masking model (mean squared error for within-channel model minus that for across-channel model = 145; p = 5.8000e-04); note that the within-channel model was simulated by replacing the CMR circuit in Figure 2B with an envelope extraction step, as in Viswanathan et al., 2022. Because the across-channel speech-intelligibility model (Figure 2B) accounts for both within-channel masking effects as well as across-channel temporal-coherence processing, this result suggests that some of the decrements in behavioral speech-in-noise performance that occur with simulated CI listening (especially current spread; rightmost plot in Figure 5) may be due to poorer across-channel temporal-coherence-based segregation of speech from background noise.

**Figure 5.**
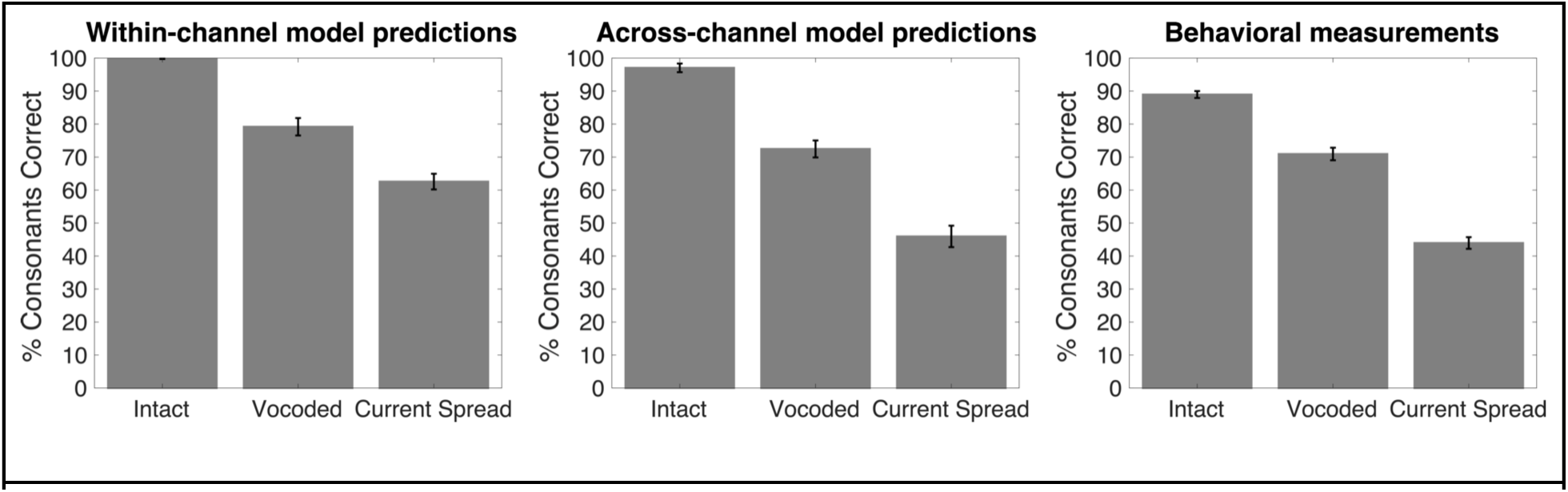
Speech intelligibility in speech-shaped noise as a function of simulated CI vocoding and current spread. The first (leftmost) plot shows predictions from a model of purely within-channel masking (mean and standard error across stimulus repetitions). The second (middle) plot shows predictions from the across-channel temporal-coherence-based source-segregation model (Figure 2B; mean and standard error across stimulus repetitions). The third (rightmost) plot shows behavioral measurements (mean and standard error across N=48 samples).

### 4.3. Simulated OHC damage degrades temporal-coherence processing

Figure 6 shows d’ estimates and CMR predictions for Intact stimuli at a fixed OFC level (83 dB SPL, which ensured target audibility at all SNRs and degrees of OHC damage) as a function of degree of OHC damage [simulated by varying the C_OHC_ parameter from 1 (normal hearing) to 0 (complete dysfunction)]. Figure 7 shows d’ estimates and CMR predictions for Intact stimuli at a fixed OFC loudness as a function of degree of OHC damage. The CMR predicted by the temporal-coherence model decreases with increasing OHC damage under both equal-SPL (Figure 6) and equal-loudness (Figure 7) conditions. Furthermore, model AN-fiber threshold tuning curves (Figure 3A) at 3225 Hz CF (i.e., the CF at which we derived CMR predictions) show that such OHC damage broadens frequency tuning, as expected. Together, these results suggest that reduction in spectral resolution from OHC damage may degrade across-channel temporal-coherence-based source segregation (of which CMR is a correlate) even for clearly audible stimuli, including those delivered through hearing-aid amplification.

**Figure 6.**
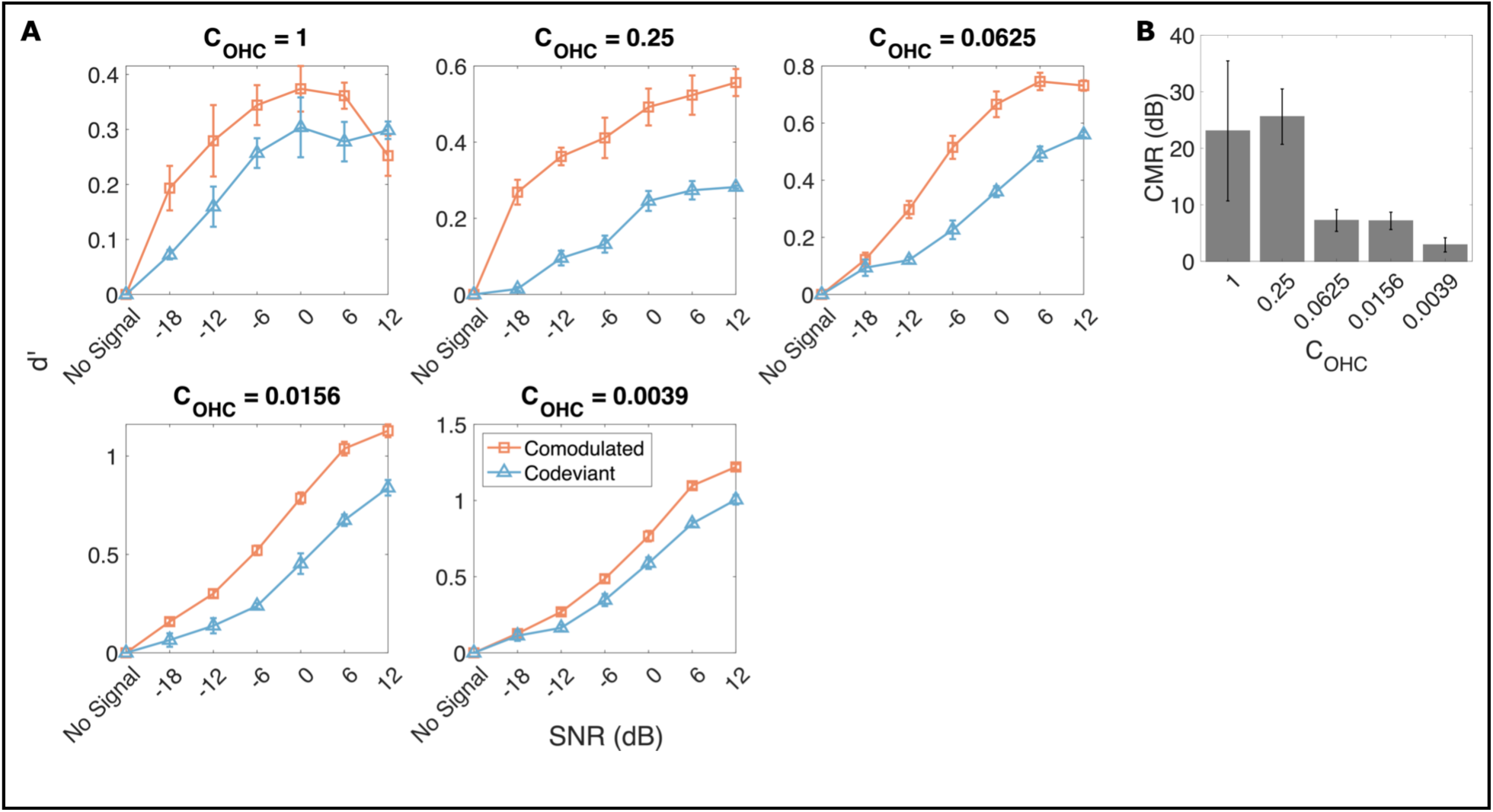
CMR predictions at a fixed OFC level (83 dB SPL) as a function of degree of simulated OHC damage [C_OHC_ varied from 1 (normal) to 0 (complete OHC dysfunction)]. The sensation level (SL) of the OFC was 63 dB at C_OHC_=1, 43 dB at C_OHC_=0.25, 28 dB at C_OHC_=0.0625, 18 dB at C_OHC_=0.0156, and 18 dB at C_OHC_=0.0039 (derived from Figure 3A). Panel A shows estimated d’ values (mean and standard error across stimulus repetitions) from the across-channel temporal-coherence-based source-segregation model (Figure 2A) for different SNRs and CMR conditions (Comodulated, Codeviant). Panel B shows CMR predictions from the temporal-coherence model (mean and standard error across stimulus repetitions), calculated as the mean difference in SNR threshold between Codeviant and Comodulated conditions across the d’ values predicted by the model.

**Figure 7.**
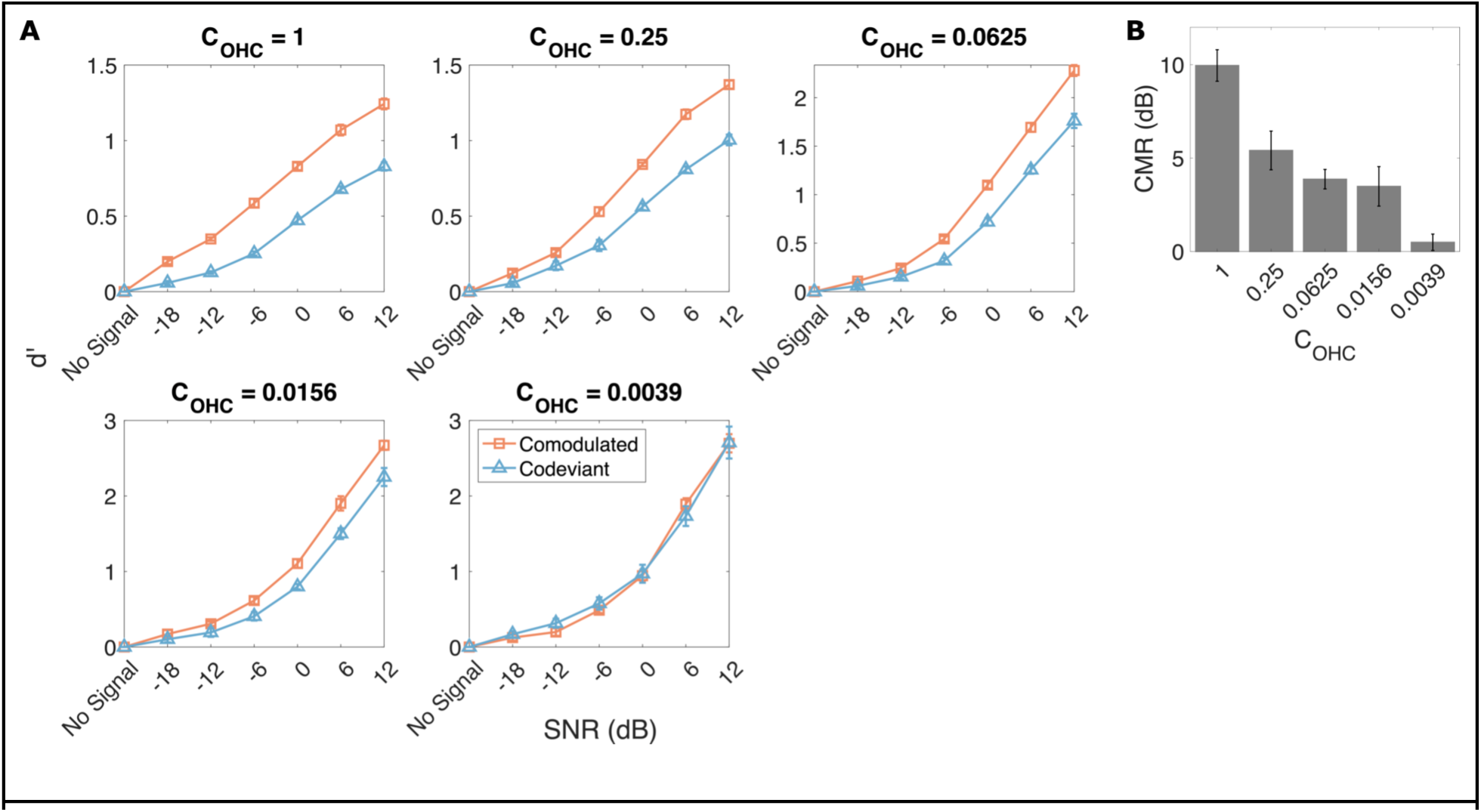
CMR predictions at a fixed OFC loudness as a function of degree of simulated OHC damage [C_OHC_ varied from 1 (normal) to 0 (complete OHC dysfunction)]. The sound pressure level (SPL) of the OFC was 48 dB at C_OHC_=1, 57 dB at C_OHC_=0.25, 64 dB at C_OHC_=0.0625, 69 dB at C_OHC_=0.0156, and 69 dB at C_OHC_=0.0039. Panel A shows d’ estimates (mean and standard error across stimulus repetitions) from the across-channel temporal-coherence-based source-segregation model (Figure 2A) for different SNRs and CMR conditions (Comodulated, Codeviant). Panel B shows CMR predictions from the temporal-coherence model (mean and standard error across stimulus repetitions), calculated as the mean difference in SNR threshold between Codeviant and Comodulated conditions across the d’ values predicted by the model.

## 5. Discussion

Using physiologically plausible computational modeling and behavioral experiments, we show that simulated CI current spread and SNHL (here, OHC damage) each adversely impact across-channel temporal-coherence-based source segregation and in turn speech-in-noise outcomes. Spectral resolution is reduced both in CI/electric hearing (especially from current spread) and in acoustic hearing with SNHL (from broadened tuning; Figure 3A). Such spectral smearing decreases sparsity (and increases across-channel correlation) in the frequency representation of different sound sources. This in turn increases the likelihood of both within-channel masking of the target by a competing sound as well as across-channel masking via grouping of temporally coherent target and masker components. Our findings underscore the importance of good peripheral frequency resolution for successful segregation of a target sound source from a distractor, like speech from background noise, and help explain why spectral smearing increases susceptibility to noise (Hall et al., 1988; Ter Keurs et al., 1992, 1993; Baer and Moore, 1993, 1994; Fu et al., 1998; Nelson et al., 2003; Stickney et al., 2004; Fu and Nogaki, 2005; Oxenham and Kreft, 2014). Note that although the vocoding filters we used for CI processing (Table 1) are slightly broader than the psychophysical tuning curves of normal-hearing listeners (Glasberg and Moore, 1990), we do not observe any significant CMR deficits for Vocoded compared to Intact stimuli; this contrasts with the large impact that simulated current spread has on CMR (Figure 4).

Our findings are consistent with the observation that CMR, a correlate of across-channel temporal-coherence processing, is smaller both in CI users (Ihlefeld et al., 2012; Zirn et al., 2013; Pierzycki and Seeber, 2014) and in hearing-impaired listeners using acoustic hearing (Hall et al., 1988; Moore et al., 1993; Ernst et al., 2010), compared to normal-hearing listeners. In CI users, experiments manipulating the degree of current spread suggest that current focusing strategies like multipolar stimulation have the potential to reduce spread of excitation relative to monopolar stimulation (Carlyon and Goehring, 2021). Our results suggest that future work on current focusing should explore strategies to improve across-channel temporal-coherence-based segregation in CI users, perhaps using specific measures like CMR in addition to overall speech-in-noise performance. Along the same lines, future work should assess whether decreased frequency selectivity in acoustic hearing with SNHL covaries with CMR measurements across individuals (Hall et al., 1988; Grose and Hall, 1996).

Although the AN model used in this study (Bruce et al., 2018) captures broadening of tuning with OHC damage, it does not capture distorted tonotopy (see Figure 3A tuning curves; Parida and Heinz, 2022b). Distorted tonotopy refers to a disruption in the mapping between acoustic frequency and cochlear place, which is caused by noise-induced hearing loss, and is often associated with greater sensitivity of a cochlear place to frequencies below its CF than to the CF itself (Henry et al., 2016, 2019). Distorted tonotopy has been suggested to be prevalent and perceptually relevant in human listeners even with only moderate hearing loss (Gruhlke et al., 2012; Kafi et al., 2022), and studies using animal models have shown that distorted tonotopy severely degrades natural speech encoding in noise, causing pathological over-representation of low-frequency sound information and background noise in the affected cochlear channels (Parida and Heinz, 2022a). This over-representation is in turn expected to impact segregation based on temporal coherence. However, because the effects of distorted tonotopy are not captured by the AN model we used in the current study (note that distorted tonotopy was captured in a previous version of this line of AN models; Heinz and Henry, 2013), the CMR predictions from our source segregation model may in fact underestimate the full impact that SNHL has on temporal-coherence-based source segregation. Future studies should be designed to specifically probe the impact of distorted tonotopy on across-channel temporal-coherence processing.

Using the modulation detection interference (MDI) paradigm, Yost and Sheft (1989) found that the detection of a modulated target tone was impaired by the presence of a masker tone with the same modulation rate as the target, even when target and masker were well separated in carrier frequency. Moreover, the modulation-depth threshold for target detection (which indicates the degree of across-channel modulation masking) is greatest when the remote masker and target are modulated in phase (Hall and Grose, 1991), in line with theories of grouping by synchrony or temporal coherence (Bregman, 1994; Elhilali et al., 2009). Surprisingly, some prior studies have reported similar MDI thresholds for normal-hearing and hearing-impaired listeners (Grose and Hall, 1994; Bacon and Opie, 2002; Sek et al., 2015). At first glance, the results of the present study, which suggest that reduced spectral resolution impacts CMR, seem at odds with these prior reports on MDI in SNHL. This discrepancy may be explained in part by the fact that some of the prior MDI studies used a non-zero phase difference between the target and masker modulations (Bacon and Opie, 2002; Sek et al., 2015), which complicates interpretation of the relationship between SNHL and across-channel temporal-coherence processing (the focus of the current study). Moreover, performance in the MDI paradigm depends on modulation-depth sensitivity/coding, which can be better in individuals with hearing loss (Schlittenlacher and Moore, 2016; Zhong et al., 2014); this also complicates interpretation. Future experiments should be designed to elucidate the precise mechanisms underlying the differential impact of hearing loss on CMR and MDI.

Our across-channel temporal-coherence-based source-segregation model is based on computations known to exist in CN (Pressnitzer et al., 2001). In this sense, it differs from other temporal-coherence-based models proposed in prior work, which are somewhat more phenomenological in nature (Elhilali et al., 2009; Christiansen et al., 2014). Although basilar membrane suppression may also contribute to perceptual CMR effects (Ernst and Verhey, 2006), little to no CMR was predicted at the output of the AN model that we used in the current study (Bruce et al., 2018) for normal hearing (see Figure 4B in Viswanathan et al., 2022). We do not model aspects of temporal-coherence processing that may exist in higher auditory stations (e.g., like the cortex; Shamma et al., 2011). Despite this, our model predictions match behaviorally measured variations in CMR and speech intelligibility in noise across the different vocoding conditions.

In an N-AFC behavioral task, the d’ statistic (effect size) is algebraically related to proportion trials correct (Green and Swets, 1966). Because our computational model does not capture all aspects of human hearing, the d’ estimated from our model cannot be expected to match the behavioral d’ and thus cannot be directly related to the behavioral proportion-correct scores. For instance, the model uses average statistics over the entire stimulus duration to estimate d’ whereas the average human subject performing tone detection in noise may not necessarily use all available data. Thus, rather than using the behavioral proportion-correct criterion (66%; Figure 4) to derive a model d’ threshold criterion, or using an arbitrary d’ threshold criterion choice, we averaged CMR predictions across all d’ values predicted by the model.

We aimed to restore loudness rather than SL in our OHC-damage simulations because under equal-SL conditions loudness recruitment is greater in SNHL compared to normal hearing (Moore, 1995). Our approach is also motivated by hearing-aid fitting procedures, which use a loudness model when calculating the necessary prescriptive amplification (Keidser et al., 2011). Despite our use of amplification to restore stimulus loudness in our OHC-damage simulations, smaller CMR is predicted with SNHL (Figure 7). This result suggests that even when using hearing aids, listeners with SNHL may experience degraded temporal-coherence processing. This degradation in turn may contribute to the perceived lack of benefit of current hearing aids (Chung, 2004; McCormack and Fortnum, 2013; Lesica, 2018).

Our model uses only medium spontaneous rate AN fibers and does not include low or high spontaneous rate fibers. This is because we wished to avoid floor and saturation effects in the AN model output firing rate, which can occur with low or high spontaneous rate fibers given our choice of stimulus levels. Note, however, that our choice of medium spontaneous rate fibers does not limit the generalizability of our results. This is because in the AN model that we used in this study, the fibers with different spontaneous rates mainly differ in their operating range of levels but are otherwise similar. Exploratory simulations with low and high spontaneous rate fibers showed similar trends to the medium spontaneous rate fiber but with floor or saturation effects for some stimulus levels (data not shown).

## Acknowledgments

This study was supported by grants from the National Institutes of Health [Grant Nos. F32DC020649 (to V.V.), R01DC019126 (to B.G.S.-C.), and R01DC009838 (to M.G.H.)]. The authors also thank Hari M. Bharadwaj for providing MATLAB code to implement linear amplification based on the Moore and Glasberg (1998, 2004) loudness model to restore audibility and loudness in hearing loss.

## Author Declarations

The authors declare no competing financial interests.

## Data Availability

The datasets used in the current study are available from V.V. on reasonable request.

